# Capturing domain-level detail of molecular interactions in the IMEx Consortium binding domain data set

**DOI:** 10.64898/2026.10.15.759378

**Authors:** Alessia Pinto, Prisca Lo Surdo, Juan Jose Medina Reyes, Lou Bergogne, Carla Padilla Franzotti, Marta Iannuccelli, Simona Panni, Kalpana Panneerselvam, Sylvie Ricard Blum, Eliot Ragueneau, Sandra Orchard, Henning Hermjakob, Andreas Zanzoni, Luana Licata

## Abstract

The machinery of the cell depends on physical interactions, between proteins themselves, and between proteins and other biomolecules. The detailed characterisation of these interactions, including the identification of the precise sequence regions mediating binding events, is essential to understand both normal cell biology and disease mechanisms. Protein binding domains are specific regions of a protein sequence that mediate direct contact with a partner and represent a critical layer of functional annotation, bridging sequence, structure, and cellular function. Here we present the IMEx binding domain dataset, a large-scale, manually curated open-access set of molecular interactions assembled over more than two decades by the International Molecular Exchange (IMEx) Consortium. The dataset compiles experimentally derived binding region annotations from peer-reviewed literature, systematically linked to protein-protein interaction (PPI) records and mapped to standardised protein sequences in UniProtKB. We describe the curation model, data format, and content of the dataset, provide a descriptive and comparative analysis, and demonstrate the utility of the dataset as a gold standard for benchmarking interface prediction and variant effect annotation methods. Its integration with genomic variations further offers a route to elucidate the mechanistic basis of disease-associated mutations. The datasets are freely available through IntAct in the dedicated PSI-MI FeatureTAB format at https://ftp.ebi.ac.uk/pub/databases/intact/current/psimitab/features/ and are updated and enhanced with every release.

## INTRODUCTION

Proteins act through networks of physical interactions with other proteins, nucleic acids, carbohydrates, lipids and small molecules, which underpins signal transduction, gene regulation, enzymatic activity, and structural organisation [1]. Characterising these networks is therefore essential to understand how cells respond to stimuli, how developmental programmes are executed, and how disruptions of interaction networks lead to disease [2,3]. Many interactions involved in signal transduction are weak but highly specific, often mediated by binding domains that recognise short linear peptides, which allows functional complexes to assemble and disassemble rapidly in response to stimuli [4]. Making effective use of PPI information requires knowing not only which proteins interact, but also through which regions. The specific segment of a protein sequence that is necessary and/or sufficient to mediate a binding event is referred to as a binding region. Binding regions range from well-characterised modular interaction domains, such as SH2 (IPR000980), SH3 (IPR001452), PDZ (IPR001478), and WD40 (IPR001680) domains [4], to interaction surfaces defined by short linear motifs (SLiMs) or intrinsically disordered regions [5]. The precise mapping of binding regions enables the identification of the molecular determinants of interaction specificity, supports the structural modelling of protein complexes, and allows the assessment of how sequence variants may disrupt or alter interaction interfaces [6,7].

The International Molecular Exchange (IMEx) Consortium was established in 2005 to provide a single, non-redundant dataset of experimentally validated molecular interactions curated to a common, internationally agreed standard [8]. Members currently includes IntAct [9], MINT [10], DIP [11], UniProt [12] and MatrixDB [13], which curate a broad range of molecular interaction data with foci on areas such as extracellular matrix biology, innate immunity, host-pathogen interactions and human disease. Curation is centralised in IntAct, allowing member resources to focus on expert annotation rather than infrastructure maintenance.

Data are distributed in community formats (PSI-MI XML2.5, XML3.0 [14] and MITAB) under a CC-BY 4.0 licence, in line with FAIR principles [15]. IMEx is recognised as an ELIXIR Core Data Resource [16] and a Global Core Biodata Resource. To date it has released 1,787,152 binary interactions curated to full IMEx standards, plus approximately 100,000 legacy records [17].

The IMEx curation model captures every interaction as an individual experimental evidence record linked to a specific publication, including the host organism and cell line, the experiment and participant detection methods, the interaction type, and the full details of the molecular constructs used, including protein isoforms, post-translational modifications (PTMs), affinity tags, and binding regions data. All proteins are mapped to their corresponding sequences in UniProtKB [12], enabling coordinate-level feature annotations that are synchronised with UniProtKB updates [17]. Annotation relies on controlled vocabulary (CV) terms from the HUPO-PSI Molecular Interactions [18].

A protein binding region is any portion of a polypeptide that mediates interaction with another molecule, ranging from a short linear motif to a modular binding domain. In the IMEx curation model, binding region data is captured in three main forms: the ‘sufficient binding region’ (MI:0442), which defines the minimum sequence sufficient to mediate a detectable interaction; the ‘necessary binding region’ (MI:0429), which identifies residues or segments required for binding; and the ‘binding-associated region’ (MI:0117), a broader term used when experimental evidence supports involvement in binding but does not allow precise delimitation of the contact surface. These annotations are typically derived from studies in which deletion mutants, truncation constructs, or defined amino acid sequences are tested for binding [17].

Binding domains are important for several reasons. First, they define the molecular logic of interaction networks: proteins may use distinct domains to interact with different partners, enabling combinatorial and context-dependent signalling. Second, they are frequent targets of post-translational modifications that promote or disrupt specific interactions depending on the resulting modification state of the protein [19]. Third, many disease-associated variants cluster at or near interaction interfaces, suggesting that disruption of specific binding events is a key disease mechanism [7,20,21]. Fourth, small molecule inhibitors can be designed to block specific protein-protein interaction interfaces, making binding domains relevant to drug discovery [22].

Several computational resources predict protein binding domains and interaction interfaces, each with a distinct scope, methodology, and data types. SMART (Simple Modular Architecture Research Tool) annotates modular protein architecture, systematically identifying and classifying signalling and extracellular domains [23]. The 3DID (Three-Dimensional Interacting Domains) database provides structural templates for domain-domain interactions observed in three-dimensional protein structures [24]. Interactome3D offers structural models of protein interactions and enables the mapping of interface residues based on structural data [25]. The ELM (Eukaryotic Linear Motif) resource focuses specifically on short linear motifs mediating regulatory interactions [26]. SLiMPred and related tools use sequence-based approaches to predict putative interaction motifs [27]. SLiMSuite offers a complementary toolkit for the discovery and evaluation of short linear motifs in protein sequences [28]. However, none of them systematically captures binding regions identified and confirmed experimentally by a wide range of methods.

Understanding how genomic variation affects protein function is central to human genetics and precision medicine [29]. Non-synonymous single nucleotide variants (nsSNVs) and other amino acid-altering variants are of particular interest, as they can disrupt protein folding, PTM sites, or interaction interfaces. Several computational tools predict the functional impact of such variants, including SIFT [30], PolyPhen-2 [31], FoldX [32] and AlphaMissense [33], but their accuracy depends critically on the quality and size of the datasets used for training and benchmarking.

Binding domain data provides a uniquely informative layer for variant effect analysis. Variants within known interaction interfaces are more likely to affect binding, as shown both for disease-associated mutations [7,20] and for engineered variants in model systems [34]. The IMEx mutations dataset [34] which currently (2026_08) contains >86,000 experimentally tested mutations and their effects on molecular interactions has proven valuable for benchmarking and for mechanistic interpretation of disease variants.

The IMEx Consortium [8] is now making available a large-scale, manually curated dataset of experimentally defined binding regions directly derived from literature evidence, systematically linked to the corresponding full interaction record, and mapped to standardised protein sequences updated with each UniProtKB release. We describe the generation of the data, analyse its content and its overlap with genomic variation data. Complementing the mutations dataset, the IMEx binding domain dataset links a variant not just to a protein but to a specific, experimentally validated binding interface.

## RESULT AND DISCUSSION

### 2.1 Dataset curation and content

The IMEx binding domain dataset is the result of more than 20 years of expert biocuration activity by the IMEx Consortium databases. Binding regions data are captured as biological features in the IntAct curation platform, following PSI-MI controlled vocabulary (CV) terms: ‘sufficient binding region’ (MI:0442), ‘necessary binding region’ (MI:0429), ‘direct binding region (MI:1125)’, ‘binding-associated region’ (MI:0117). Whenever a binding site corresponds to a known structural domain or repeated element, a cross-reference to the corresponding InterPro [35] entry is added at the feature level. This linking step is performed manually by curators, when the domain identity is reported by the original authors; here, we complement it with an automated mapping pipeline that assigns an InterPro domain to a binding region when the two approximately coincide (at least 80% overlap in both directions; see Section 2.2 and Methods). Together, these provide a direct connection between experimentally defined binding regions and the broader landscape of protein domain families.

The full dataset currently contains 232,649 binding region annotations distributed across 177,021 interaction records with at least one binding region annotated. The vast majority correspond to ‘sufficient binding region’ annotations (179,203 in total across all organisms), which represent experimental evidence from fragment or deletion construct studies. ‘Binding-associated region’ annotations number 48,124, while ‘necessary binding region’ annotations, which require more stringent experimental demonstration, account for 5,311 entries. The dataset spans a wide taxonomic range, covering 1,038 species, although the distribution is strongly weighted towards well-studied eukaryotic model organisms (Table 1). Human proteins account for the largest share of annotations, with 133,425 sufficient binding regions, 19,558 binding-associated regions, and 2,571 necessary binding region entries in the current dataset. Other well-represented organisms include *Mus musculus* (12,297 total binding region annotations), *Rattus norvegicus* (3,873), *Drosophila melanogaster* (10,028), *Saccharomyces cerevisiae* (9,776), *Caenorhabditis elegans* (4,794), and *Escherichia coli* (1,378) (Figure 1, Table 1).

**Table 1.** Counts and within-species percentages of each binding region type (from the Molecular Interaction ontology, MI), computed at the interaction level for each of the seven species. The blue colour intensity encodes the within-species percentage only.

| Binding domain regions |  |  |  |
| --- | --- | --- | --- |
| Species | Binding domain type | Count | % (within species) |
| *H. sapiens* | Sufficient binding region (MI:0442) | 133,425 | 85.8 |
| *H. sapiens* | Binding-associated region (MI:0117) | 19,558 | 12.6 |
| *H. sapiens* | Necessary binding region (MI:0429) | 2,571 | 1.7 |
| *M. musculus* | Sufficient binding region (MI:0442) | 7,954 | 64.7 |
| *M. musculus* | Binding-associated region (MI:0117) | 3,827 | 31.1 |
| *M. musculus* | Necessary binding region (MI:0429) | 516 | 4.2 |
| *S. cerevisiae* | Sufficient binding region (MI:0442) | 3,498 | 35.8 |
| *S. cerevisiae* | Binding-associated region (MI:0117) | 5,962 | 61.0 |
| *S. cerevisiae* | Necessary binding region (MI:0429) | 316 | 3.2 |
| *R. norvegicus* | Sufficient binding region (MI:0442) | 1,743 | 45.0 |
| *R. norvegicus* | Binding-associated region (MI:0117) | 1,992 | 51.4 |
| *R. norvegicus* | Necessary binding region (MI:0429) | 138 | 3.6 |
| *D. melanogaster* | Sufficient binding region (MI:0442) | 5,712 | 57.0 |
| *D. melanogaster* | Binding-associated region (MI:0117) | 4,223 | 42.1 |
| *D. melanogaster* | Necessary binding region (MI:0429) | 93 | 0.9 |
| *E. coli* | Sufficient binding region (MI:0442) | 875 | 63.5 |
| *E. coli* | Binding-associated region (MI:0117) | 391 | 28.4 |
| *E. coli* | Necessary binding region (MI:0429) | 112 | 8.1 |
| *C. elegans* | Sufficient binding region (MI:0442) | 4,558 | 95.0 |
| *C. elegans* | Binding-associated region (MI:0117) | 179 | 3.7 |
| *C. elegans* | Necessary binding region (MI:0429) | 57 | 1.2 |

**Figure 1.**
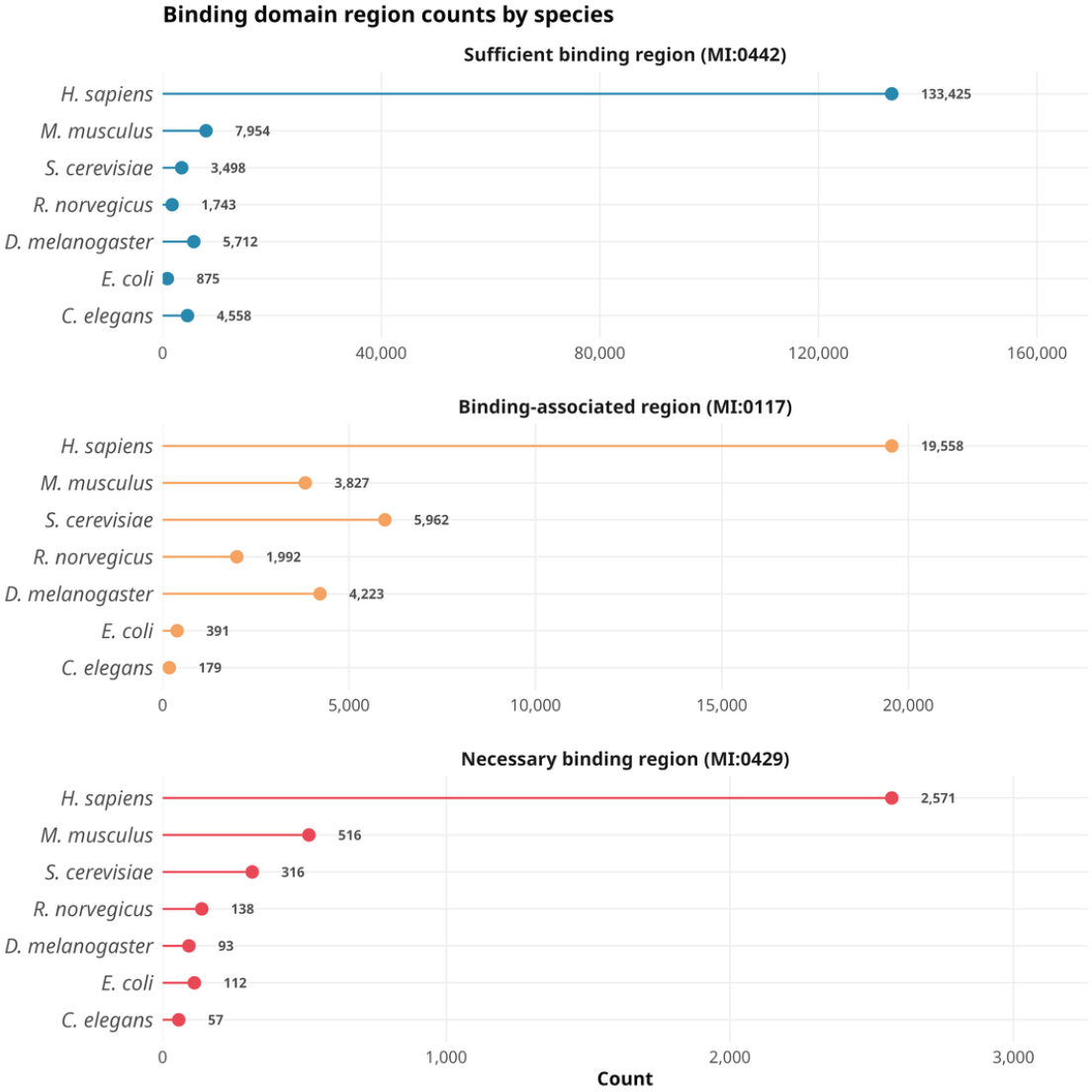
Lollipop plot of the counts of each binding region type (MI ontology) at the interaction level, shown per species and split by region type.

The distribution of annotations per publication follows a highly skewed pattern consistent with that observed in the IMEx mutations dataset [34]: the large majority of publications contribute fewer than ten annotations each, reflecting the dominance of small-scale, hypothesis-driven studies in the literature. A small number of publications contribute large numbers of annotations, corresponding to systematic domain-mapping studies or high-throughput interaction screens that report binding regions for many protein pairs simultaneously (Figure 2A).

**Figure 2.**
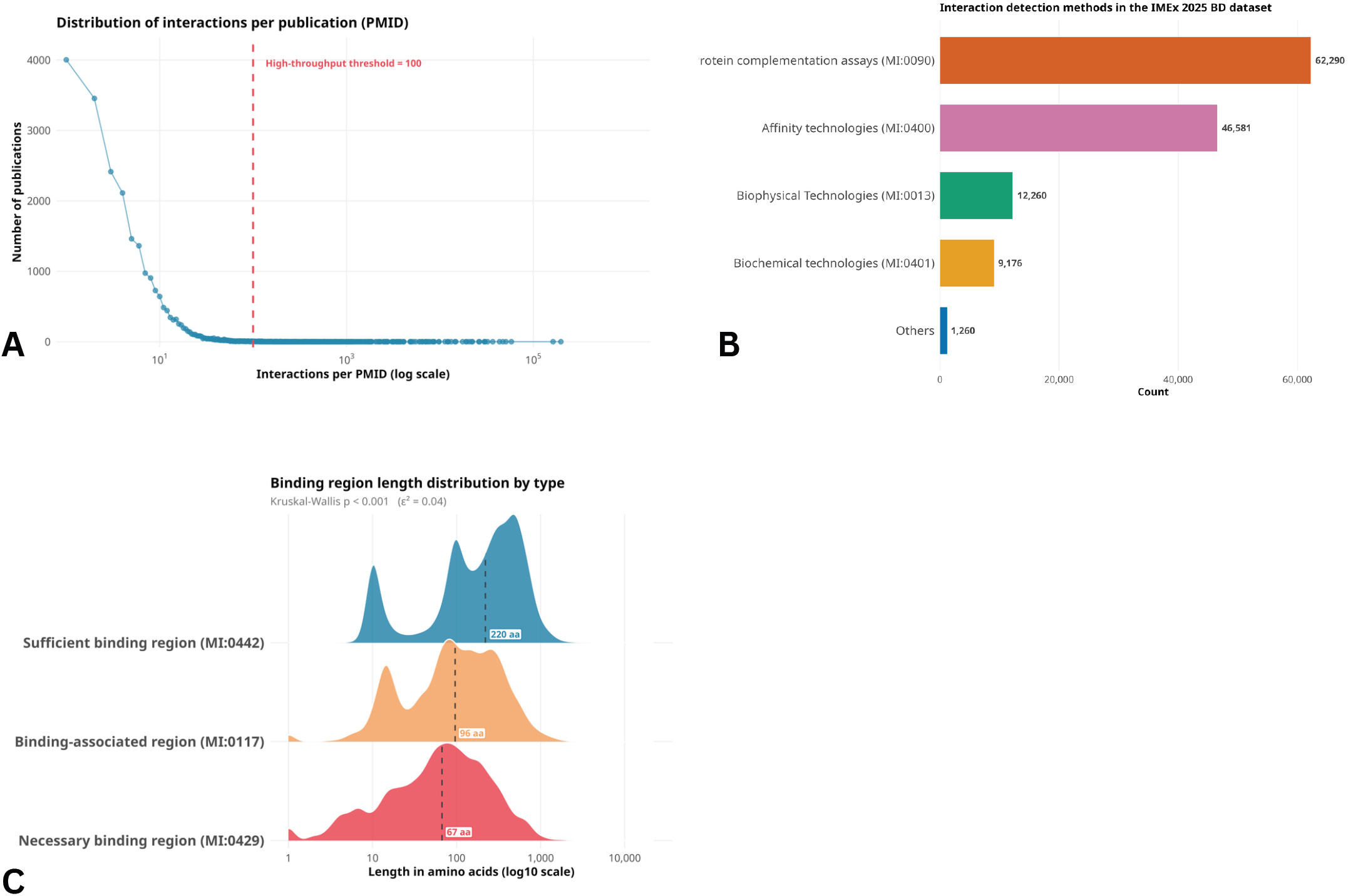
A) Distribution of interactions per publication (PMID) in the IMEx 2025 release, used to define the low-throughput subset (cut-off: 100 interactions per publication). Most publications report only one or a few interactions, and the frequency drops sharply as the number of interactions per publication increases. The x-axis (interactions per PMID) is log-transformed; the y-axis shows the number of publications. B) Barplot of the number of binding-region-annotated interactions by interaction detection method (MI-standardised), showing the most frequently used methods. C) Length distributions of the three binding region types annotated in the IMEx 2025 human dataset (length on a log scale). Within each distribution, the dashed line and label mark the median. A non-parametric Kruskal–Wallis test was performed; the p-value and the effect size (epsilon-squared) are reported.

In terms of interaction detection methods, protein complementation assays, primarily variations of the yeast two-hybrid system, account for the largest share of binding region annotations, followed by affinity chromatography-based techniques such as co-immunoprecipitation and pull-down assays. Biophysical methods, including surface plasmon resonance and isothermal titration calorimetry, contribute a smaller but methodologically distinct subset of annotations (Figure 2B). Kinetic data, such as dissociation rates (kds) and equilibrium dissociation constants (KDs) reflecting the affinity of interactions are often generated by such experiments and are captured as part of the annotation process. This distribution reflects the types of experimental approaches most commonly used to delineate interaction interfaces in the primary literature.

The three binding region types also differ in their length distributions (Figure 2C). Sufficient binding regions (median of 220 amino acids), which derive from fragment and deletion constructs, and binding-associated regions (median of 96 amino acids) span a broad range of lengths, whereas necessary binding regions (median of 67 amino acids), which pinpoint the residues strictly required for binding, tend to be more sharply delimited. A non-parametric Kruskal–Wallis test confirms that the three length distributions differ significantly, in agreement with the distinct experimental resolution captured by each annotation type (p-value < 0.001). Restricting the analysis to the low-throughput subset, in which each publication reports at most 100 interactions, alters the relative percentage composition of the three binding region types (Supplementary Figure 1). The relative composition of the dataset therefore partly reflects the contribution of a small number of high-throughput studies, which should be considered when the dataset is used for benchmarking.

### 2.2 Mapping to InterPro domain families

In IMEx, an InterPro cross-reference is added to a binding region only when the domain is explicitly identified in the original publication, which limits coverage to a small fraction of binding regions. To place the whole dataset in the context of known protein domain architecture, we therefore developed a mapping pipeline that identifies all InterPro domains overlapping each binding region and quantifies the extent of the overlap in both directions. For the annotation of the dataset, only domains that overlap above 80% of the binding region are retained.

We first checked which InterPro domains are preferentially located within annotated binding regions rather than elsewhere along the protein sequence. For each domain, Fisher’s exact test compared its instances coinciding with a binding region against those not overlapping any binding region (FDR < 0.05). Among the domains that coincide with an experimentally defined binding region, the most over-represented are well-known protein interaction modules (Figure 3A). They include extracellular ligand and receptor domains, such as the tumour necrosis factor, C1q, PDGF/VEGF and chemokine domains and the recognition domains of immune receptors, as well as intracellular binding modules such as 14-3-3 and the SH3 domain of endophilin-A, and the dimerisation domain of basic-leucine zipper transcription factors (IPR004827), which shows the largest number of instances and the strongest statistical support. These results indicate that, where a binding region matches a single InterPro domain, the domain itself typically corresponds to the interaction interface identified experimentally. At the same time, the prominence of extensively studied domain families may partly reflect their over-representation in the literature rather than a biological property alone.

**Figure 3.**
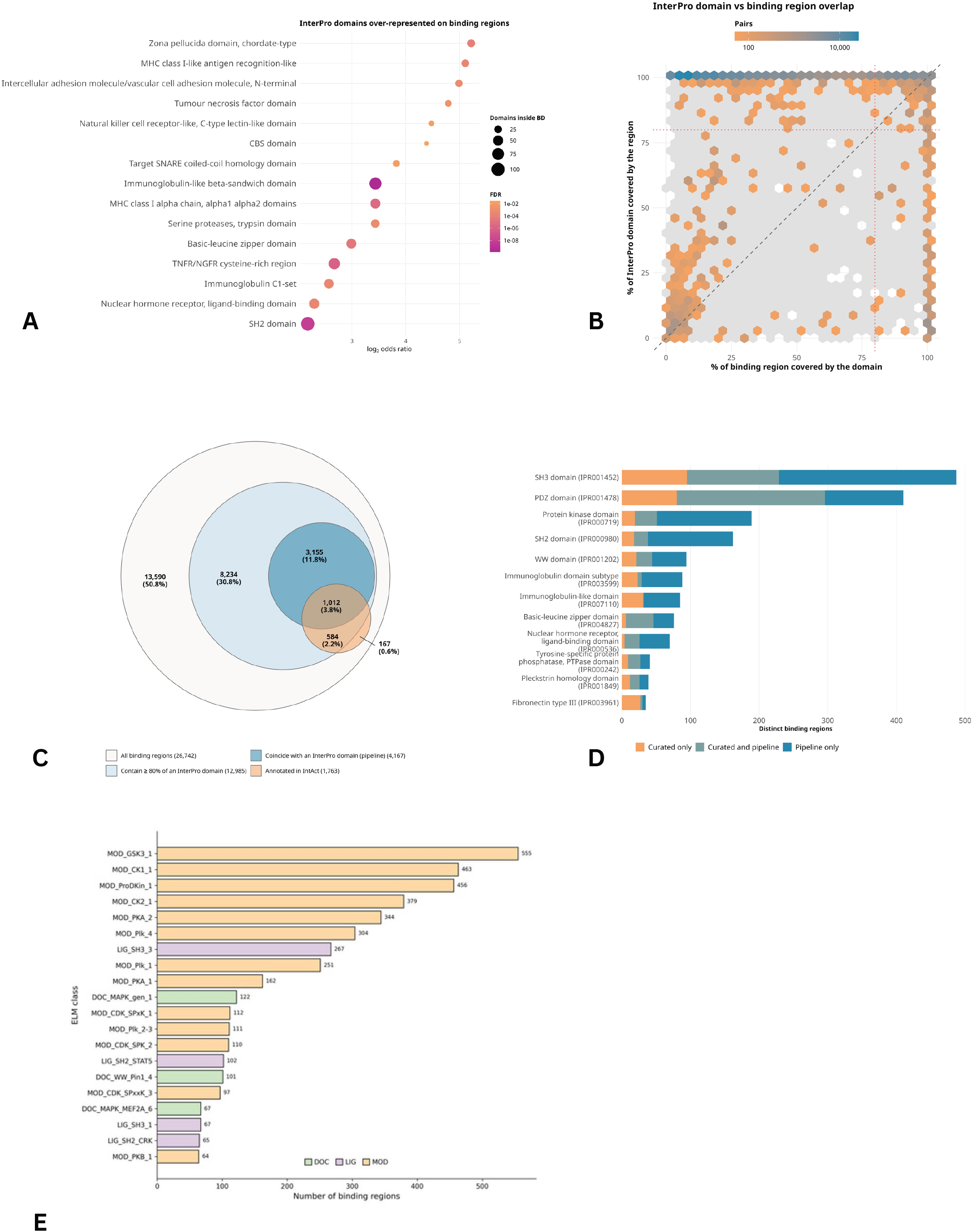
A) InterPro domains over-represented within binding regions. For each InterPro domain, regions having at least 80% overlap in both directions were compared with instances not overlapping any binding region, using a one-sided Fisher’s exact test (BH FDR < 0.05). The fifteen domains with the highest odds ratio are shown. Dot colour encodes the FDR and dot size the number of domain instances coinciding with a binding region. B) Hexbin density of the overlap between percentage of feature and percentage of domain. Colour intensity encodes the number of overlapping (binding region, InterPro domain) pairs per cell; cells with fewer than 50 pairs are shown in grey. C) Euler diagram of the human distinct human binding regions (grey). Light blue, binding regions containing at least 80% of an InterPro domain; blue, the subset of regions having at least 80% overlap in both directions; orange, regions carrying a curator-assigned InterPro domain entry. (D) For the twelve InterPro domains most frequently assigned by curators, number of distinct binding regions carrying the domain according to curation only (orange), to both curation and the pipeline with the same InterPro entry (grey-green), or to the pipeline only (blue). E) Distribution of non redundant ELM classes (i.e. motifs) instances mapped to binding regions. ELM classes are colored according to class type. DOC: docking sites; LIG: ligand binding sites; MOD: post-translational modification sites. The overlap threshold is set to 0.8.

Consistently, an over-representation analysis of the proteins whose binding regions coincide exactly with a single InterPro domain returns functional annotation terms centred on signalling, actin cytoskeleton and Rho-GTPase dynamics, and membrane trafficking (Supplementary Figure 3), showing that this domain-level pattern is also reflected at the level of biological functions and pathways.

We then measured how InterPro domains and binding regions overlap across all region-domain pairs identified by the pipeline (Figure 3B). For each pair, we calculated two values: the fraction of the domain covered by the binding region, and the fraction of the binding region occupied by the domain. Most pairs show 100% domain coverage, while the fraction of the binding region occupied by the domain varies widely. In other words, InterPro domains are usually fully contained within larger binding regions rather than matching them exactly, suggesting that binding interfaces often extend beyond the boundaries of a single annotated domain.

A binding region that contains a domain is not necessarily one in which that domain mediates the interaction. For this reason, the pipeline assigns an InterPro domain to a binding region only when the two approximately coincide, that is, when at least 80% of the domain falls within the region and at least 80% of the region falls within the domain.

Of the 26,742 distinct human binding regions, 12,985 (48.6%) contain at least 80% of one or more InterPro domains. In most of these cases (8,818; 33.0% of all regions), however, the region extends well beyond the domain. Only 4,167 regions (15.6%) meet the coincidence criterion and are assigned an InterPro domain (Figure 3C).

We then compared these assignments with curator annotations. Curators had assigned an InterPro domain to 1,763 regions (6.6%). The pipeline recovers 1,012 of them (57.4%), and in 976 cases (96.4%) it assigns the same InterPro entry. Of the 751 curated regions not recovered, 584 contain the annotated domain but extend well beyond it, while 167 do not contain at least 80% of any InterPro domain. Finally, the pipeline assigns a domain to 3,155 regions (11.8%) that had no curator-assigned domain (Figure 3C, D).

Together, these analyses show that the IMEx binding domain dataset covers well-characterised interaction domains involved in signalling, cytoskeletal dynamics and membrane trafficking, and that binding regions frequently extend beyond the annotated domain. They also show that systematic mapping to InterPro can extend domain annotation to regions not previously annotated by curators.

### 2.4 Mapping to ELM linear motif classes

In addition to structured domains, a significant fraction of protein interactions is mediated by short linear motifs (SLiMs): short, rapidly evolving sequence elements found predominantly in intrinsically disordered regions that bind to structured domains on partner proteins [5]. To characterise this component of the IMEx binding domain dataset, we mapped ELM (Eukaryotic Linear Motif) classes [26] to all binding region annotations using a coordinate overlap threshold of 0.8.

The mapping reveals a substantial number of IMEx binding features that correspond to known ELM motif classes (Figure 3E), extending the functional annotation of the IMEx dataset beyond globular domains. Motif classes involved in protein binding, docking sites, and phosphorylation-dependent interactions are among the most represented (Figure 3E, Supplementary Table 1). For the latter, we checked whether the required phospho-residue has been previously detected as such in phospho-proteomics experiments [36]. We indeed observed that most of the phosphorylation-dependent motif instances include an experimentally identified phospho-residue (Supplementary Figure 4). This finding highlights an important dimension of the dataset: binding regions captured by IMEx curators frequently correspond to putative functional sequence elements that are not described as classical InterPro domains, and whose experimental characterisation in the context of physical interactions adds value to purely sequence-based motif resources. The combination of InterPro and ELM mappings thus provides a more complete picture of the sequence determinants of protein interactions represented in the dataset.

**Figure 4.**
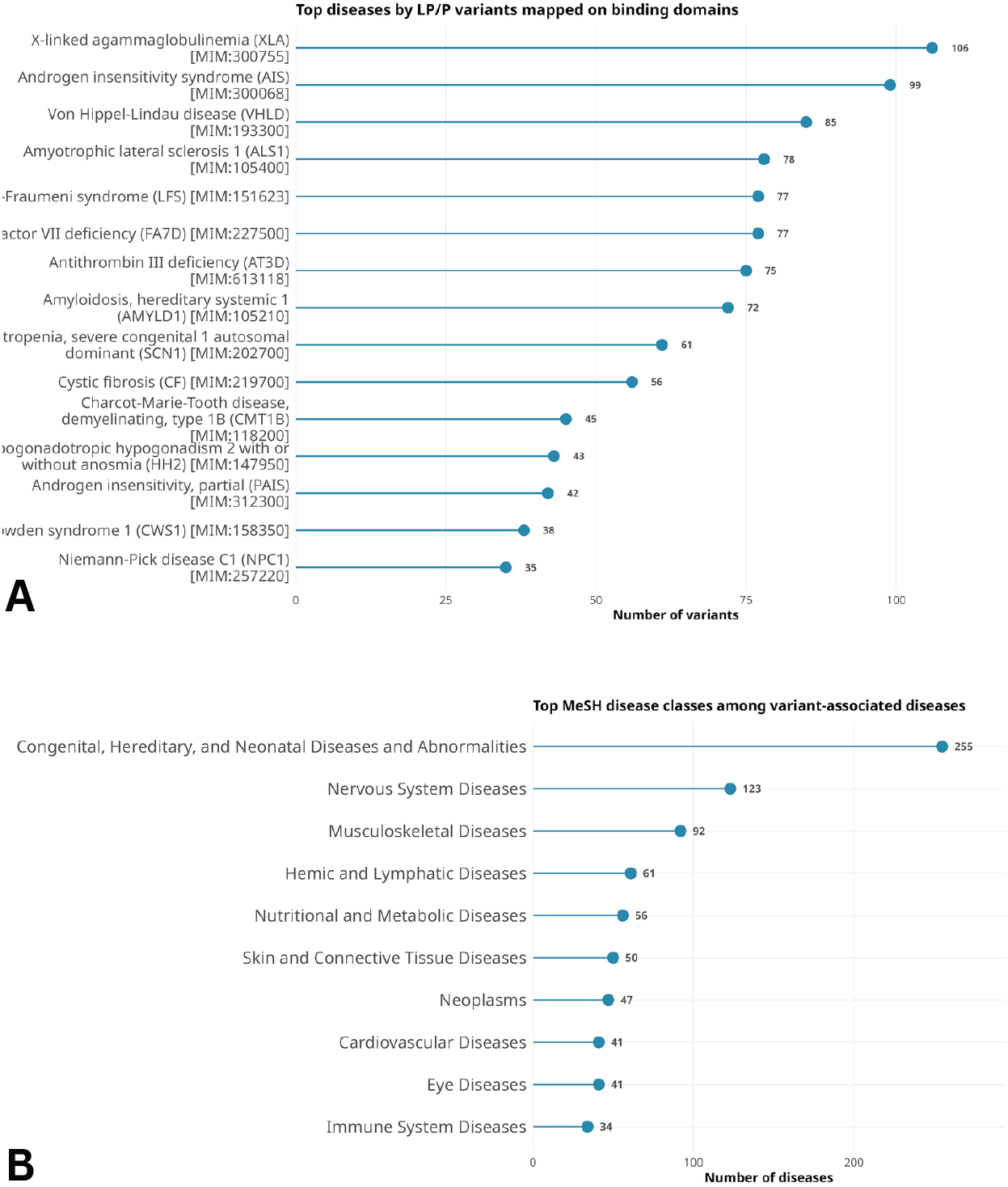
A) The fifteen diseases with the most LP/P variants falling within the overlap between binding regions and InterPro domains. Each variant is counted once, regardless of the number of binding regions, interactions or InterPro domains in which it falls. B) MeSH classification of the diseases associated with LP/P variants located within binding regions overlapping an InterPro domain, showing the number of diseases in each top-level MeSH class. Only diseases with at least one MeSH class are shown. The predominance of congenital, hereditary and neonatal diseases is consistent with most of these variants being associated with Mendelian disorders.

### 2.5 Disease variant analysis: LP/P variants and variants of uncertain significance

A key application of the IMEx binding domain dataset is the interpretation of genomic variants, particularly in a clinical context. To explore this application, we mapped human disease-associated variants from UniProtKB [12] onto the binding region annotations in the human IMEx dataset. Variants were classified as likely pathogenic or pathogenic (LP/P) based on their annotation status in UniProtKB, which integrates manual curation and cross-references from ClinVar [37] and other variant resources.

Of the 31,949 unique LP/P variants annotated in UniProtKB, 19,817 fall on proteins carrying at least one IMEx binding region. Of these, 11,722 (59.2%), of 1,261 proteins, lie within an experimentally defined binding region. Restricting the analysis to binding regions that approximately coincide with an InterPro domain retains 3,122 variants on 353 proteins, which can be assigned both to a specific interaction interface and to the domain that mediates it. A large substantial number of known pathogenic variants therefore lies within regions supported by experimental binding data. Rather than being linked only to the protein as a whole, these variants can be placed within a specific, experimentally defined binding interface, offering a starting point for the mechanistic interpretation of their effect.

The diseases with the most LP/P variants within the binding region-domain overlaps are X-linked agammaglobulinemia and androgen insensitivity syndrome (Figure 4A). More generally, these diseases fall into several functional groups: disorders of signalling proteins and hormone receptors, such as X-linked agammaglobulinemia, complete and partial androgen insensitivity and hypogonadotropic hypogonadism; inherited cancer predisposition syndromes caused by tumour suppressors, including von Hippel–Lindau disease, Li-Fraumeni syndrome and Cowden syndrome; coagulation disorders, such as factor VII and antithrombin III deficiencies; and diseases associated with protein misfolding or neurodegeneration, including amyotrophic lateral sclerosis 1, hereditary systemic amyloidosis and Charcot–Marie–Tooth disease type 1B. In all these cases, pathogenic variants fall within domains that coincide with experimentally defined interaction interfaces, consistent with the disruption of specific protein-protein interactions as a possible mechanism of pathogenicity. We also classified the diseases associated with the mapped LP/P variants by MeSH (Medical Subject Headings) category. The distribution was broad, with the largest numbers of diseases in Congenital, Hereditary, and Neonatal Diseases and Abnormalities class (Figure 4B).

We extended this analysis to variants of uncertain significance (VUS), the category for which mechanistic information is most needed. VUS within binding region-domain overlaps are dominated by cancer-associated entries (Supplementary Figure 2). Most of these describe sporadic tumours or individual cancer samples rather than defined Mendelian diseases; a smaller number are associated with inherited conditions such as open-angle glaucoma, long QT syndrome 1 and Loeys–Dietz syndrome. This is consistent with observations from the IMEx mutations dataset [34] and with structural analyses showing that cancer variants preferentially target protein-protein interaction interfaces [21]. For VUS in particular, knowing that a variant falls within an experimentally characterised binding region provides a concrete starting point for experimental follow-up and for the development of computational models of variant effect. Together, these results show that the IMEx binding domain dataset can support variant interpretation. The represented disease associations span immunodeficiencies, endocrine and coagulation disorders, cancer predisposition syndromes and neurodegenerative diseases and multiple cancer types. This breadth reflects the biological coverage of the underlying IMEx interaction data, and suggests that the dataset can be applied across a wide range of clinical genomics contexts.

### 2.6 Data access and visualisation

The complete IMEx Binding Domain dataset is freely available for download from the IntAct FTP server (https://ftp.ebi.ac.uk/pub/databases/intact/current/psimitab/features/) under a CC-BY 4.0 licence, and is updated with each IntAct release. In addition to the standard PSI-MI XML and MITAB representations, the binding-domain annotations are distributed in the PSI-MI FeatureTAB format, a tab-delimited data-interchange format developed by the HUPO-PSI Molecular Interactions working group specifically to capture IMEx feature annotations at the sequence level [34]. Unlike PSI-MI MITAB and XML, which are designed to exchange experiment-based interaction records, FeatureTAB is intended for specialist use cases that require the systematic extraction of sequence features, such as variants, post-translational modifications and binding regions, together with their molecular and experimental context.

Individual binding regions can also be examined within the context of a specific interaction using ComplexViewer [38], an open-source viewer embedded in IntAct and Complex Portal (https://github.com/MICommunity/ComplexViewer). For each interactor, ComplexViewer now integrates DisProt [39], AlphaFold [40], ELM [26] and InterPro [35] annotations as cross-referenced tracks alongside the curated binding region, each linked to its source database, allowing a curator or researcher to assess, case by case, the structural context flanking an annotated binding region (see Supplementary Figure 5 for an example on the TP53–MDM2 interaction).

## METHODS

### Data sources

The August 2025 IMEx release was used in this study, downloaded from the IntAct portal (https://www.ebi.ac.uk/intact/interactomes). In addition, the November 2024 release was used for the ELM class mapping.

### Software and Package used

Binding region extraction, plotting, and interaction with the InterPro database (including querying the InterPro API and merging the retrieved annotations) were performed with standard Python libraries (pandas, numpy, re, ast, requests). Figures and ORA analyses were produced in R (libraries: ggplot2 [41], clusterProfiler [42], ReactomePA [43], gt, ggridges).

### IMEx Dataset Analysis

IMEx dataset contains 232,649 total binding region annotations. Exploratory analyses were conducted on the 2025 IMEx dataset for seven species: *Homo sapiens* (taxon 9606), *Saccharomyces cerevisiae* (taxon 559292), *Drosophila melanogaster* (taxon 7227), *Caenorhabditis elegans* (taxon 6239), *Mus musculus* (taxon 10090), *Escherichia coli* K12 (taxon 83333), and *Rattus norvegicus* (taxon 10116). Using custom Python scripts, binding region annotations were extracted for each target species. An interaction was retained when at least one of the two interactors belonged to the target species; interactions in which neither interactor belonged to the target species were discarded. The species-specific datasets therefore also include cross-species interactions in which only one partner is a target-species protein. Binding region annotations were counted only on the interactors belonging to the target species, so that each annotation is attributed to the species of the protein involved (Table 1). A sub-dataset restricted to interactions annotated in low-throughput experiments was also derived for each species dataset, by removing publications (PubMed identifiers) reporting more than 100 interactions (Supplementary figure 1).

Three binding region types from the Molecular Interaction (MI) vocabulary were quantified: sufficient binding region (MI:0442), necessary binding region (MI:0429), and binding-associated region (MI:0117). For each species we report the number of binding region annotations at the interaction level, together with the number of distinct proteins carrying each annotation type, expressed also as a percentage of all annotated proteins in that species. Direct binding region annotations were retained in the dataset but excluded from the counts, as they represented a negligible number of entries (11 entries).

Binding region length was defined using length coordinates (length = end − start + 1). The three length distributions (by binding region type) were compared with a non-parametric Kruskal–Wallis test on a log scale; given the large sample size, the effect size (epsilon-squared) and the group medians are reported alongside the p-values.

### InterPro Domain Mapping

InterPro domains were mapped onto the binding region annotations of the August IMEx 2025 human dataset using the InterPro API. InterPro annotations were retrieved for all 9,242 accessions carrying at least one binding region (1,564 isoforms; processed chains were not queried): 9,206 returned InterPro entries and 7,746 at least one entry of type ‘domain’, the only type used for the mapping. Where available, the specific isoform annotated in the IMEx dataset was queried; otherwise the canonical sequence was used, and isoforms carrying their own InterPro annotations were treated as distinct entries.

Binding regions without explicit numeric boundaries (8,081 ranges, e.g. undetermined or terminal positions) were discarded, leaving 132,735 interactor-level binding region records on 8,990 accessions (7,816 distinct genes once isoforms are collapsed to their canonical sequence). Of these, 7,493 accessions (6,474 genes; 1,340 isoforms mapped independently) had at least one InterPro domain, and 6,448 (5,623 genes) had at least one domain overlapping a binding region.

The overlap was defined as the number of shared amino acids between the two regions (overlap = max(0, min(ends) − max(starts) + 1)), and two metrics were computed for each region-domain pair:

Percentage of domain = (overlap / InterPro domain length) × 100

Percentage of feature = (overlap / binding region length) × 100,

where percentage of domain is the fraction of the InterPro domain covered by the binding region, and percentage of feature is the fraction of the binding region covered by the InterPro domain. In the IMEx binding domain dataset, pipeline-derived InterPro cross-references are added to a binding region only when the two approximately coincide, i.e. with at least 80% of the domain covered by the region and at least 80% of the region covered by the domain. A high-confidence subset was derived from the complete mapping, restricted to entries reaching 100% for both metrics (318 unique proteins, 307 genes).

Titin (Q8WZ42), annotated with a single binding region spanning residues 2787–34350 in a high-throughput study, was excluded from all InterPro-based analyses.

The comparison between curated and pipeline-derived annotations was performed on distinct binding regions, defined by protein accession, binding region type and boundaries, so that a region annotated in several interactions was counted once. A region was considered curator-annotated when its own IntAct feature carried an InterPro identifier. Over-representation of InterPro domains within binding regions was assessed on all InterPro domain instances (entries of type ‘domain’) annotated on human proteins carrying at least one binding region with numeric boundaries. Each domain instance was classified as inside when it approximately coincided with at least one binding region of the same protein (at least 80% of the domain covered by the region and at least 80% of the region covered by the domain, on the same region), as outside when it did not overlap any binding region, and as partial otherwise. Partial instances were excluded, so that domains coinciding with a binding region were compared with domains lying entirely outside binding regions. For each InterPro entry with at least three instances, a 2×2 contingency table (inside/outside × this domain/all other domains) was tested with a one-sided Fisher’s exact test for enrichment. Odds ratios were reported as effect size, and p-values were adjusted with the Benjamini–Hochberg procedure (FDR < 0.05). Domains with at least five instances were considered for display (Figure 3A).

Over-representation analysis (ORA) was performed on the high confidence subset of proteins whose binding regions completely overlap a single InterPro domain (for both Percentage of Feature and Percentage of Domain at 100), using as background all proteins with at least one InterPro domain overlapping a binding region; isoforms were collapsed to their canonical accession. The analysis was run against Gene Ontology (Biological Process and Molecular Function) [44], KEGG [45], and Reactome [46], with p-values adjusted by the Benjamini–Hochberg procedure (FDR < 0.05).

### Mapping ELM classes in Binding Domain

The IMEx 2024 dataset (downloaded on 2024-11-19) has been filtered to recover reviewed human-human PPIs only. All interactors sequences have been gathered from UniProtKB [12] rest API. Short linear motifs were detected using the SLiMProb tool, via the *mimic*INT workflow [47], and the motif definitions from the Eukaryotic Linear Motif (ELM) database (version 1.4, 2024) [26]. Motif search was limited to predicted disordered regions identified by IUPred [48, 49], with both short and long modalities, and with a propensity score threshold equal to 0.2 [28]. Phosphorylation data was downloaded from the PhosphoSitePlus resource [36] (June 2023).

### Mapping variants in Binding Domains and Disease Categories

Human variants curated from literature reports were downloaded from UniProtKB (humsavar.txt, release 2025_03), with their associated pathogenicity classifications (LP/P, likely pathogenic/pathogenic; LB/B, likely benign/benign; US, uncertain significance) and associated disease annotations from OMIM [50] and cancer samples. Variant positions, numbered on the canonical sequence, were mapped only onto binding regions annotated on canonical proteins. From the LP/P and VUS variants, two subsets were extracted: variants located within an IMEx binding region, and variants located within the intersection between a binding region and an InterPro domain, restricted to region–domain pairs that approximately coincide (at least 80% reciprocal coverage, as above) and excluding titin. Variants were deduplicated by protein and amino acid change; mapping rates are reported relative to the variants located on proteins carrying at least one binding region. For the disease-level analysis, each variant was counted once per disease, regardless of the number of binding regions, interactions or InterPro domains in which it falls. Diseases associated with the variants within binding region–domain overlaps were assigned a MeSH disease class through a two-step cascade: MeSH classes were taken directly from DisGeNET version 26.2 [51] (curated entries only), and for diseases not classified there, the Unified Medical Language System (UMLS CUI) provided by DisGeNET was used to retrieve the corresponding top-level MeSH code from Wikidata [52]. Only top-level MeSH disease categories were retained. Because a disease can map to more than one MeSH category, classes are not mutually exclusive; diseases with no MeSH class were labelled “Unclassified”. Overall, 55.4% of variant-associated diseases received at least one MeSH class.

## CONCLUSIONS

The IMEx binding domain dataset collects 232,649 experimentally derived binding region annotations across 177,021 interaction records and 1,038 species, curated from the literature over more than two decades and mapped to UniProtKB sequences. Each annotation is linked to the full experimental record in which it was observed, including constructs, detection methods and publication. The dataset is distributed under a CC-BY 4.0 licence in PSI-MI XML, MITAB and FeatureTAB formats and is updated with every IntAct release.

Beyond describing the dataset, this work provides two resources. First, we release an extended version of the IMEx binding domain dataset in which each binding region, annotated for a specific interactor in a specific interaction, is linked to the InterPro domains with which it approximately coincides. These links combine the cross-references assigned by curators with those derived by our mapping pipeline, which extends domain annotation to 3,155 human binding regions that previously lacked one, broadening the range of applications for which the dataset can be used. Second, we make available the mapping pipeline itself (see Code Availability), which retrieves InterPro annotations for any set of IMEx binding regions, retains only the domains that match the binding region according to user-defined coverage criteria, and identifies the individual residues falling within each binding region–domain overlap, allowing, for instance, sequence variants to be placed within a specific, experimentally supported interaction interface.

Our analyses show that binding regions frequently extend beyond annotated domains, that many contain short linear motifs, and that 11,722 pathogenic or likely pathogenic variants fall within experimentally defined binding regions.

The dataset reflects the composition of the literature: well-studied proteins and domain families, as well as a small number of high-throughput studies, are over-represented, and this should be taken into account when the data are used for training or benchmarking. Within these limits, the dataset can support the development and evaluation of interface and motif predictors and the prioritisation of variants, particularly VUS, for experimental follow-up.

The IMEx Consortium continues to curate binding regions, and future IntAct releases will include the InterPro cross-references derived by the pipeline.

## AUTHOR CONTRIBUTIONS

Alessia Pinto: Formal Analysis, Investigation, Methodology, Visualisation, Writing – Original Draft; Prisca Lo Surdo: Formal Analysis, Methodology, Visualisation, Writing – Original Draft Preparation; Juan Jose Medina Reyes: Investigation; Lou Bergogne: Formal Analysis, Investigation, Methodology, Visualisation; Carla Padilla Franzotti: Visualisation; Kalpana Panneerselvam: Data Curation; Marta Iannuccelli: Data Curation; Simona Panni: Data Curation, Writing—review & editing; Sylvie Ricard Blum: Data Curation, Writing—review & editing; Eliot Ragueneau: Investigation; Sandra Orchard: Data Curation, Writing—review & editing; Henning Hermjakob: Investigation, Supervision, Writing – Review & Editing; Andreas Zanzoni: Investigation, Supervision, Writing – Review & Editing; Luana Licata: Conceptualization, Data Curation, Formal Analysis, Funding Acquisition, Investigation, Methodology, Supervision, Writing – Original Draft Preparation, Writing – Review & Editing.

## ACKNOWLEDGEMENTS

The authors would like to thank all IMEx Consortium curators who generated the binding region annotations: P. Porras, B. Meldal, M Duesbury, L. Perfetto, J. Khadake, D. Thorneycroft, K. van Roey, S. Abbani, L. Salwinski, M. Pellegrini, B. Roechert, A. Bridge, M. G. Ammari, F. McCarthy, F. Broackes-Carter, N. H. Campbell, A. N. Melidoni, M. Rodriguez-Lopez, R. C. Lovering, S. Jagannathan, C. Chen, D. J. Lynn, U. Mahadevan, and A. Raghunath

## REFERENCES

1. Barabási, A.-L. & Oltvai, Z.N. Network biology: understanding the cell’s functional organization. Nat. Rev. Genet. 5, 101–113 (2004).

2. Luck, K., Sheynkman, G.M., Zhang, I. & Vidal, M. Proteome-scale human interactomics. Trends Biochem. Sci. 42, 342–354 (2017).

3. Correia, F.B. et al. Handling noise in protein interaction networks. Biomed. Res. Int. 2019, 8984248 (2019).

4. Pawson, T. & Nash, P. Assembly of cell regulatory systems through protein interaction domains. Science 300, 445–452 (2003).

5. Van Roey, K. et al. Short linear motifs: ubiquitous and functionally diverse protein interaction modules directing cell regulation. Chem. Rev. 114, 6733–6778 (2014).

6. Fragoza, R. et al. Extensive disruption of protein interactions by genetic variants across the allele frequency spectrum in human populations. Nat. Commun. 10, 4141 (2019).

7. Sahni, N. et al. Widespread macromolecular interaction perturbations in human genetic disorders. Cell 161, 647–660 (2015).

8. Orchard, S. et al. Protein interaction data curation: the International Molecular Exchange (IMEx) consortium. Nat. Methods 9, 345–350 (2012).

9. Del Toro, N. et al. The IntAct database: efficient access to fine-grained molecular interaction data. Nucleic Acids Res. 50, D648–D653 (2022).

10. Licata, L. et al. MINT, the molecular interaction database: 2012 update. Nucleic Acids Res. 40, D857–D861 (2012).

11. Salwinski, L. et al. The Database of Interacting Proteins: 2004 update. Nucleic Acids Res. 32, D449–D451 (2004).

12. UniProt Consortium. UniProt: the Universal Protein Knowledgebase in 2025. Nucleic Acids Res. 53, D609–D617 (2025).

13. Samarasinghe, K.W. et al. MatrixDB 2024: an increased coverage of extracellular matrix interactions, a new Network Explorer and a new web interface. Nucleic Acids Res. 53, D1677–D1682 (2025).

14. Sivade Dumousseau, M. et al. Encompassing new use cases — level 3.0 of the HUPO-PSI format for molecular interactions. BMC Bioinformatics 19, 134 (2018).

15. Wilkinson, M.D. et al. The FAIR Guiding Principles for scientific data management and stewardship. Sci. Data 3, 160018 (2016).

16. Drysdale, R. et al. The ELIXIR Core Data Resources: fundamental infrastructure for the life sciences. Bioinformatics 36, 2636–2642 (2020).

17. Porras, P. et al. Towards a unified open access dataset of molecular interactions. Nat. Commun. 11, 6144 (2020).

18. Hermjakob, H. et al. The HUPO PSI’s molecular interaction format — a community standard for the representation of protein interaction data. Nat. Biotechnol. 22, 177–183 (2004).

19. Lee, M.J. & Yaffe, M.B. Protein regulation in signal transduction. Cold Spring Harb. Perspect. Biol. 8, a005918 (2016).

20. Wang, X. et al. Three-dimensional reconstruction of protein networks provides insight into human genetic disease. Nat. Biotechnol. 30, 159–164 (2012).

21. Porta-Pardo, E. et al. A pan-cancer catalogue of cancer driver protein interaction interfaces. PLoS Comput. Biol. 11, e1004518 (2015).

22. Arkin, M.R., Tang, Y. & Wells, J.A. Small-molecule inhibitors of protein–protein interactions: progressing toward the reality. Chem. Biol. 21, 1102–1114 (2014).

23. Letunic, I. & Bork, P. SMART v10: three decades of the protein domain annotation resource. Nucleic Acids Res. 54, D499–D503 (2026).

24. Stein, A., Russell, R.B. & Aloy, P. 3DID: interacting protein domains of known three-dimensional structure. Nucleic Acids Res. 33, D413–D417 (2005).

25. Mosca, R., Céol, A. & Aloy, P. Interactome3D: adding structural details to protein networks. Nat. Methods 10, 47–53 (2013).

26. Kumar, M. et al. The Eukaryotic Linear Motif resource: 2024 update. Nucleic Acids Res. 52, D442–D455 (2024).

27. Mooney, C., Pollastri, G., Shields, D.C. & Haslam, N.J. Prediction of short linear protein binding regions. J. Mol. Biol. 415, 193–204 (2012).

28. Edwards, R.J., Paulsen, K., Aguilar Gomez, C.M. & Pérez-Bercoff, Å. Computational prediction of disordered protein motifs using SLiMSuite. Methods Mol. Biol. 2141, 37–72 (2020).

29. Claussnitzer, M. et al. A brief history of human disease genetics. Nature 577, 179–189 (2020).

30. Kumar, P., Henikoff, S. & Ng, P.C. Predicting the effects of coding non-synonymous variants on protein function using the SIFT algorithm. Nat. Protoc. 4, 1073–1081 (2009).

31. Adzhubei, I.A. et al. A method and server for predicting damaging missense mutations. Nat. Methods 7, 248–249 (2010).

32. Van Durme, J. et al. A graphical interface for the FoldX forcefield. Bioinformatics 27, 1711–1712 (2011).

33. Cheng, J. et al. Accurate proteome-wide missense variant effect prediction with AlphaMissense. Science 381, eadg7492 (2023).

34. IMEx Consortium Curators. Capturing variation impact on molecular interactions in the IMEx Consortium mutations data set. Nat. Commun. 10, 10 (2019).

35. Blum, M. et al. InterPro: the protein sequence classification resource in 2025. Nucleic Acids Res. 53, D444–D456 (2025).

36. Hornbeck, P.V. et al. 15 years of PhosphoSitePlus®: integrating post-translationally modified sites, disease variants and isoforms. Nucleic Acids Res. 47, D433–D441 (2019).

37. Landrum, M.J. et al. ClinVar: updates to support classifications of both germline and somatic variants. Nucleic Acids Res. 53, D1313–D1321 (2025).

38. Combe, C.W. et al. ComplexViewer: visualization of curated macromolecular complexes. Bioinformatics 33, 3673–3675 (2017).

39. Nugnes, M.V. et al. DisProt in 2026: enhancing intrinsically disordered proteins accessibility, deposition, and annotation. Nucleic Acids Res. 54, D383–D392 (2026).

40. Varadi, M. et al. AlphaFold Protein Structure Database: massively expanding the structural coverage of protein-sequence space with high-accuracy models. Nucleic Acids Res. 50, D439–D444 (2021).

41. Wickham, H. ggplot2: Elegant Graphics for Data Analysis (Springer, 2016).

42. Wu, T. et al. clusterProfiler 4.0: a universal enrichment tool for interpreting omics data. Innovation 2, 100141 (2021).

43. Yu, G. & He, Q.-Y. ReactomePA: an R/Bioconductor package for reactome pathway analysis and visualization. Mol. BioSyst. 12, 477–479 (2016).

44. The Gene Ontology Consortium. The Gene Ontology knowledgebase in 2023. Genetics 224, iyad031 (2023).

45. Kanehisa, M., Furumichi, M., Sato, Y., Matsuura, Y. & Ishiguro-Watanabe, M. KEGG: biological systems database as a model of the real world. Nucleic Acids Res. 53, D672–D677 (2025).

46. Milacic, M. et al. The Reactome Pathway Knowledgebase 2024. Nucleic Acids Res. 52, D672–D678 (2024).

47. Choteau, S.A. et al. mimicINT: a workflow for microbe-host protein interaction inference. F1000Research 14, 128 (2025).

48. Dosztányi, Z., Csizmók, V., Tompa, P. & Simon, I. The pairwise energy content estimated from amino acid composition discriminates between folded and intrinsically unstructured proteins. J. Mol. Biol. 347, 827–839 (2005).

49. Dosztányi, Z. Prediction of protein disorder based on IUPred. Protein Sci. 27, 331–340 (2018).

50. Amberger, J.S., Bocchini, C.A., Scott, A.F. & Hamosh, A. OMIM.org: leveraging knowledge across phenotype–gene relationships. Nucleic Acids Res. 47, D1038–D1043 (2019).

51. Piñero, J. et al. The DisGeNET knowledge platform for disease genomics: 2019 update. Nucleic Acids Res. 48, D845–D855 (2020).

52. Vrandečić, D. & Krötzsch, M. Wikidata: a free collaborative knowledgebase. Commun. ACM 57, 78–85 (2014).

