## Supplementary figures for "Capturing domain-level detail of molecular interactions in the IMEx Consortium binding domain data set"

1

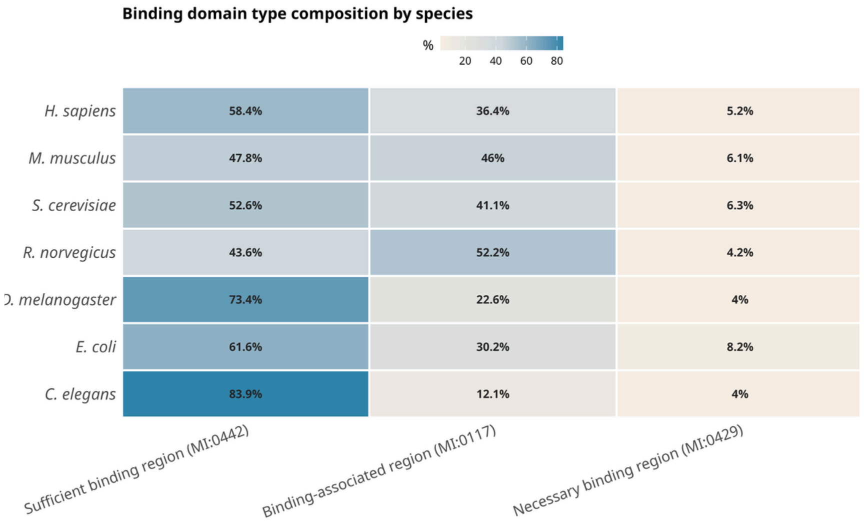


**Supplementary Figure 1.** Heatmap of the within-species composition of the three binding region types at the interaction level in the low-throughput subset; colour intensity encodes the percentage within each species-specific dataset.

2

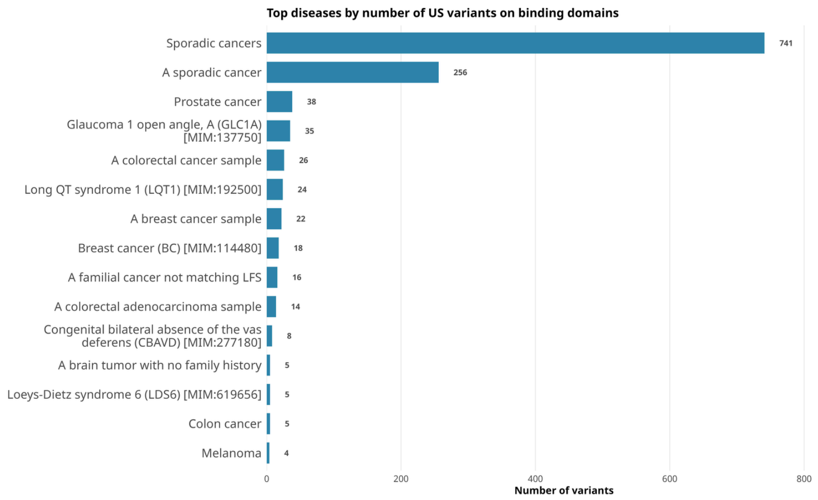


**Supplementary Figure 2.** The fifteen diseases with the most VUS falling within binding region-domain overlaps, using the same criteria as in Figure 4A. Each variant is counted once; titin is excluded. Disease labels are reported as annotated in UniProtKB humsavar; several cancer-related entries describe tumour samples rather than distinct diseases.

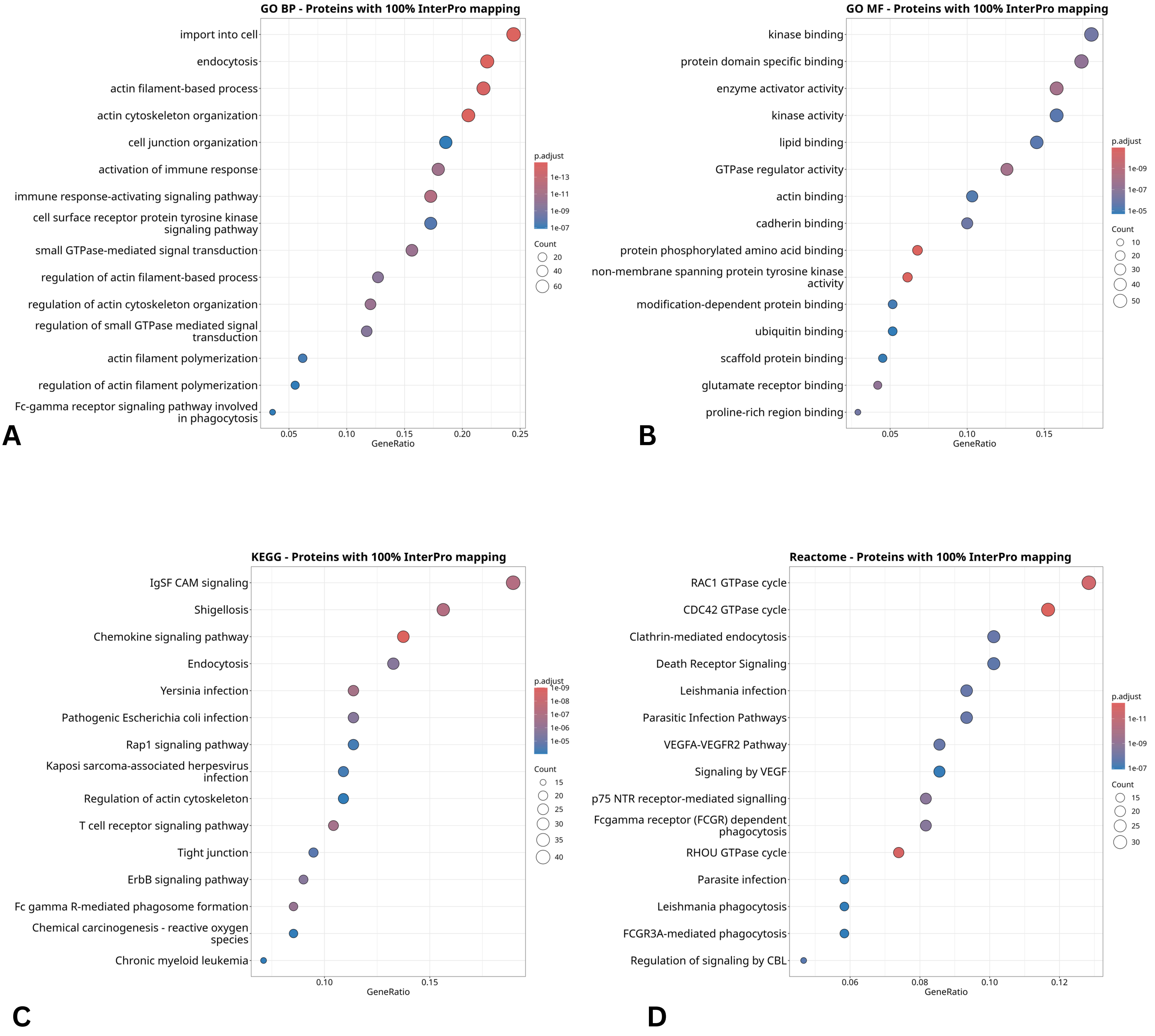


**3**

**Supplementary Figure 3.** Over-representation dot plots (top 15 terms) for the proteins with 100% overlap between the IMEx-annotated binding region and the InterPro domain, across Gene Ontology Biological Process (A), Gene Ontology Molecular Function(B), KEGG (C), and Reactome (D). Term significance (adjusted p-value) is shown by colour and gene count by dot size.

4

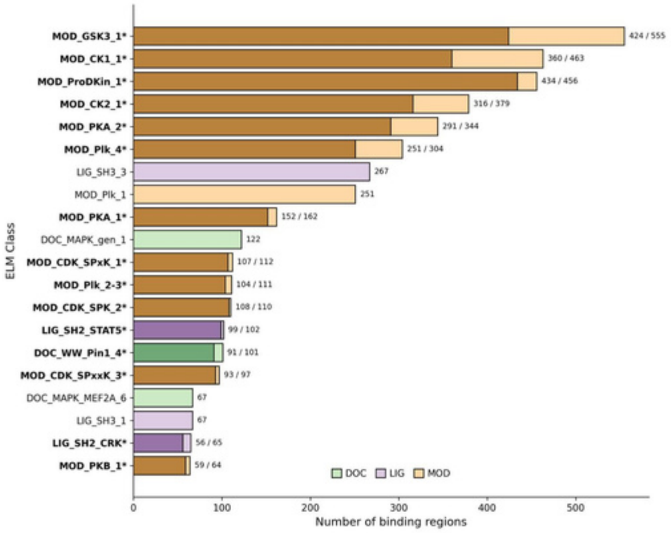


**Supplementary Figure 4.** Distribution of non redundant ELM classes (i.e. motifs) instances mapped to binding regions. ELM classes requiring a phospho-residue are reported in bold and noted with an asterisk. ELM classes are colored according to class type. Dark shades of color denote the number of instances in which the required phospho-residue has been detected as such experimentally. DOC: docking sites; LIG: ligand binding sites; MOD: post-translational modification sites. The overlap threshold is set to 0.8.

5

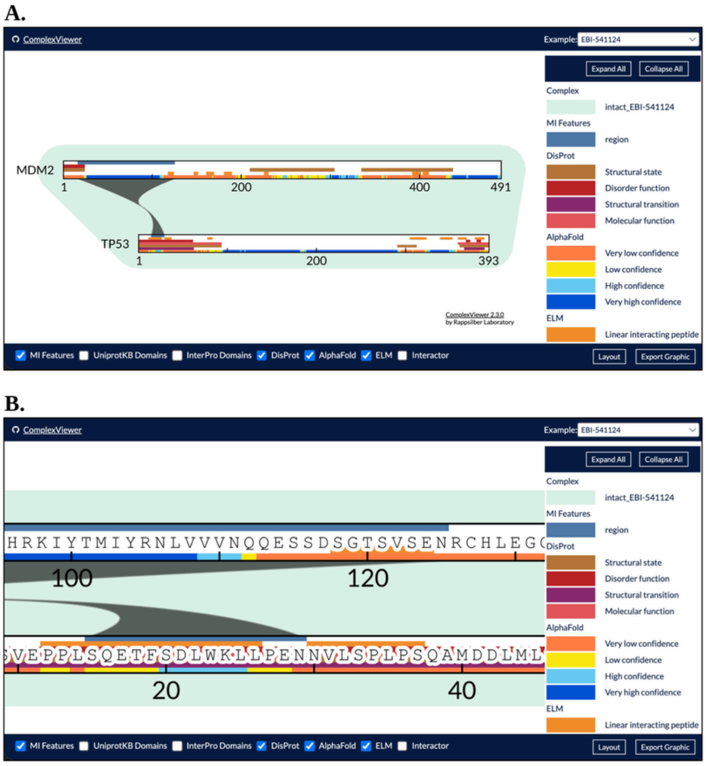


**Supplementary Figure 5.** ELM motifs within curated binding regions, visualised at the level of an individual interaction using ComplexViewer (v2.3.0) for the TP53-MDM2 interaction (IntAct EBI-541124). Panel A: the binding regions of TP53 (P04637, residues 15-29) and MDM2 (Q00987, residues 17-125) are linked in the record and shown as a shaded band. DisProt and AlphaFold tracks show the structural context of each motif; InterPro and UniProtKB domain tracks are not shown in this example. Panel B: residue-level view of the ELM motifs within the binding regions. Supplementary Table 2 provides detailed information on the motifs shown in this example.

| ELM_class | # nr instances in binding regions | # nr instances with phospho-residues |
|---|---|---|
| MOD_GSK3_1 | 555 | 424 |
| MOD_CK1_1 | 463 | 360 |
| MOD_ProDKin | 456 | 434 |
| MOD_CK2_1 | 379 | 316 |
| MOD_PKA_2 | 344 | 291 |
| MOD_Plk_4 | 304 | 251 |
| LIG_SH3_3 | 267 | 0 |
| MOD_Plk_1 | 251 | 0 |
| MOD_PKA_1 | 162 | 152 |
| DOC_MAPK_g | 122 | 0 |
| MOD_CDK_SP | 112 | 107 |
| MOD_Plk_2-3 | 111 | 104 |
| MOD_CDK_SP | 110 | 108 |
| LIG_SH2_STA | 102 | 99 |
| DOC_WW_Pin | 101 | 91 |
| MOD_CDK_SP | 97 | 93 |
| DOC_MAPK_M | 67 | 0 |
| LIG_SH3_1 | 67 | 0 |
| LIG_SH2_CRK | 65 | 56 |
| MOD_PKB_1 | 64 | 59 |
| LIG_SH2_SRC | 61 | 60 |
| LIG_SH3_2 | 49 | 0 |
| LIG_PDZ_Clas | 48 | 0 |
| MOD_DYRK1A | 42 | 42 |
| LIG_WW_1 | 38 | 0 |
| LIG_SH2_GRB | 38 | 37 |
| LIG_SH2_STA | 35 | 31 |
| MOD_LATS_1 | 27 | 27 |
| DOC_USP7_M | 27 | 0 |
| LIG_SH2_STA | 22 | 22 |
| DOC_PP2A_B | 22 | 0 |
| DEG_APCC_D | 20 | 0 |
| LIG_FHA_2 | 19 | 13 |
| DOC_ANK_TN | 18 | 0 |
| LIG_FHA_1 | 17 | 16 |
| LIG_PTB_Pho | 16 | 16 |
| LIG_PTB_Apo | 16 | 0 |
| LIG_SH3_PxxD | 16 | 0 |
| DEG_SCF_FB | 16 | 16 |
| LIG_BRCT_BR | 14 | 13 |
| DOC_MAPK_R | 14 | 0 |

**Supplementary Table 1**

| | | |
|---|---|---|
| MOD_PIKK_1 | 14 | 14 |
| DEG_MDM2_S | 14 | 0 |
| LIG_Pex14_1 | 13 | 0 |
| DOC_MAPK_N | 13 | 0 |
| TRG_ENDOCY | 12 | 0 |
| MOD_LOK_Yx | 12 | 12 |
| MOD_NEK2_1 | 11 | 10 |
| LIG_ULM_U2A | 10 | 0 |
| DOC_MAPK_H | 10 | 0 |
| DEG_SCF_TR | 9 | 9 |
| LIG_TYR_ITIM | 9 | 9 |
| LIG_TYR_ITSM | 9 | 9 |
| LIG_PTAP_UE | 9 | 0 |
| LIG_PAM2_1 | 8 | 0 |
| LIG_PDZ_Clas | 8 | 0 |
| TRG_NLS_Mo | 8 | 0 |
| LIG_SH3_4 | 7 | 0 |
| DOC_MAPK_D | 7 | 0 |
| MOD_SUMO_ | 7 | 0 |
| DEG_SPOP_S | 6 | 0 |
| LIG_PDZ_Clas | 6 | 0 |
| DEG_SCF_FB | 6 | 6 |
| LIG_GSK3_LR | 5 | 0 |
| LIG_BRCT_BR | 5 | 5 |
| LIG_BRCT_MD | 5 | 0 |
| LIG_NRBOX | 5 | 0 |
| TRG_ER_diArg | 5 | 0 |
| LIG_MAD2 | 5 | 0 |
| LIG_LYPXL_S_ | 4 | 0 |
| LIG_SxIP_EBH | 4 | 0 |
| LIG_SH3_Pxxx | 4 | 0 |
| DOC_PP4_Fxx | 4 | 0 |
| TRG_NLS_Mo | 4 | 0 |
| LIG_SH3_CIN | 4 | 0 |
| DOC_CYCLIN | 3 | 0 |
| MOD_AAK1BI | 3 | 3 |
| DEG_ODPH_V | 3 | 0 |
| DEG_Kelch_K | 3 | 0 |
| DEG_COP1_1 | 3 | 0 |
| DOC_CYCLIN | 3 | 0 |
| TRG_DiLeu_B | 3 | 0 |
| LIG_DLG_GKl | 3 | 3 |

**Supplementary Table 1**

| | | |
|---|---|---|
| LIG_MYND_1 | 3 | 0 |
| LIG_EH_1 | 3 | 0 |
| DOC_SPAK_O | 2 | 0 |
| LIG_TRAF2like | 2 | 0 |
| LIG_APCC_AB | 2 | 0 |
| DOC_CYCLIN | 2 | 0 |
| DOC_USP7_U | 2 | 0 |
| DOC_AGCK_P | 2 | 0 |
| LIG_CaM_IQ_ | 2 | 0 |
| LIG_IRFs_LxIS | 2 | 2 |
| LIG_PDZ_Wm | 2 | 0 |
| TRG_NLS_Mo | 1 | 0 |
| DOC_AGCK_P | 1 | 0 |
| TRG_DiLeu_B | 1 | 0 |
| DEG_SCF_SK | 1 | 0 |
| TRG_ER_FFAT | 1 | 1 |
| TRG_ER_FFAT | 1 | 0 |
| DEG_APCC_K | 1 | 0 |
| DEG_CRL4_C | 1 | 0 |
| LIG_MYND_3 | 1 | 0 |
| LIG_VCP_SHP | 1 | 0 |
| DOC_MAPK_J | 1 | 0 |
| LIG_CAP-Gly_ | 1 | 0 |
| LIG_PCNA_PI | 1 | 0 |
| LIG_MTR4_AIM | 1 | 0 |
| LIG_FAT_LD_1 | 1 | 0 |
| LIG_RB_pABg | 1 | 0 |
| LIG_EVH1_1 | 1 | 0 |
| LIG_Dynein_D | 1 | 0 |
| LIG_CaM_NSC | 1 | 0 |
| LIG_Actin_RP | 1 | 0 |
| DOC_PP1_RV | 1 | 0 |
| LIG_TRAF6_M | 1 | 0 |
| LIG_ANK_PxL | 1 | 0 |
| LIG_UBA3_1 | 1 | 0 |
| LIG_WRC_WI | 1 | 0 |
| MOD_CDC14 | 1 | 1 |
| DOC_PP2B_P | 1 | 0 |
| DOC_PP2B_L | 1 | 0 |
| CLV_PCSK_SK | 1 | 0 |

### Supplementary Table 1

**Supplementary Table 1.** Binding regions with overlapping ELM classes.

| Protein | UniProt | Curated binding region | ELM instance | ELM class | Start-End | Motif sequence | Motif coverage | ELM evidence (PMID) |
|---|---|---|---|---|---|---|---|---|
| TP53 | P04637 | 15–29 | ELMI000949 | MOD_PIKK_1 | 12–18 | PPLSQET | 0.57 | 9733515 |
| | | | ELMI002838 | DEG_MDM2_SWIB_1 | 19–26 | FSDLWKLL | 1.0 | 8875929 |
| MDM2 | Q00987 | 17–125 | ELMI003828 | MOD_CK1_1 | 118–124 | SGTSVSE | 1.0 | 20708156 |

### Supplementary Table 2

**Supplementary Table 2.** ELM motif instances within or overlapping the binding regions shown in Supplementary Figure 5. All instances are curated in ELM as true positives (Logic “TP”). Motif coverage is the fraction of the motif covered by the binding region. DEG_MDM2_SWIB_1 (highlighted in grey) mediates the TP53-MDM2 interaction, whereas MOD_PIKK_1 and MOD_CK1_1 are phosphorylation sites recognised by ATM/ATR and CK1δ, respectively.
